# Beyond pain: a brain-based biomarker predicts individual pain relief

**DOI:** 10.64898/2026.08.21.746216

**Authors:** Jialin Li, Balint Kincses, Katharina Schmidt, Katarina Forkmann, Lea Busch, Jaspreet Kaur, Frederik Schlitt-Nguyen, Katja Wiech, Ulrike Bingel, Tamas Spisak

**Affiliations:** Department of Neurology, Center for Translational Neuro- and Behavioral Sciences (C-TNBS), University Medicine Essen, University Duisburg-Essen, 45147 Essen, Germany; Max Planck School of Cognition, 04103 Leipzig, Germany; Oxford Centre for Integrative Neuroimaging (OxCIN), Nuffield Department of Clinical Neurosciences, University of Oxford, John Radcliffe Hospital, Oxford OX3 9DU, UK; Center for Translational Neuro- and Behavioral Sciences (C-TNBS), University Hospital Essen, Essen, Germany

## Abstract

Effective pain relief is a central goal of medical care, yet objective biomarkers of pain relief are lacking. Using task-based functional MRI and a capsaicin-induced tonic heat pain model, we experimentally elicited both pain exacerbation and pain relief within the same individuals. Existing brain-based signatures, including the Neurologic Pain Signature (NPS), reliably detected pain increases, but failed to capture pain relief. We therefore developed the PAin and RElief Signature (PARES), a multivariate brain signature trained to predict bidirectional changes in pain perception in n = 61 healthy controls. PARES robustly predicted both pain increases and relief and generalized to an independent cohort of people with chronic back pain (n = 58), who underwent the same experimental procedures. Together these findings establish a neural signature of pain relief and provide a potential biomarker for treatment stratification and analgesic development.

**One sentence summary:** A brain-based fMRI signature captures both pain exacerbation and relief, providing a candidate biomarker for analgesic development.

## INTRODUCTION

Relief from pain is a primary goal of medical treatment and a key determinant of patient wellbeing and recovery, with insufficient relief increasing the risk of transition to chronic pain (*1–3*). Yet pain relief in response to pharmacological and non-pharmacological interventions varies substantially across individuals and remains difficult to predict (*4*), posing a major challenge for clinical decision-making and analgesic development. Objective, quantifiable biomarkers of the inherently subjective experience of pain relief could help address this challenge by providing mechanistically interpretable measures of treatment effects, facilitating analgesic evaluation and development, and ultimately supporting more personalized approaches to pain medicine.

Over the past decade, interest has grown in developing brain-based biomarkers of pain that provide an objective and quantifiable readout of a fundamentally subjective experience. This effort has led to the development of multivariate brain signatures such as the Neurologic Pain Signature (NPS), which reliably tracks experimentally evoked pain intensity across individuals and paradigms (*5–7*), and the Stimulus Intensity Independent Pain Signature (SIIPS), which captures pain-related variance beyond nociceptive input, including cognitive and affective modulatory processes (*8*). Together, these signatures have provided robust markers of pain-related processes and have been validated across a wide range of experimental and clinical contexts. However, whether existing brain-based pain signatures generalize to pain relief remains an important open question. This question is particularly relevant because pain relief is unlikely to represent merely the inverse of pain amplification, but may involve partly distinct neurobiological processes (*2, 9, 10*). Consistent with opponent-process and reinforcement-learning accounts (*11*), relief from ongoing pain has been proposed to constitute a distinct affective and motivational state rather than simply the absence of nociceptive signaling (*12*). Yet existing brain-based pain signatures have primarily been developed and validated to predict pain intensity rather than reductions in ongoing pain.

Here, we addressed this gap by experimentally eliciting both pain exacerbation and pain relief within the same individuals using functional magnetic resonance imaging (fMRI) and a capsaicin-based tonic heat pain model. We first tested whether the NPS and SIIPS generalize to pain relief. While both signatures robustly detected pain increase, neither captured subjective reports of pain relief. We therefore developed the PAin and RElief Signature (PARES), a multivariate brain signature trained to predict bidirectional changes in pain perception. PARES accurately predicted both pain increase and relief and generalized to an independent cohort of people with chronic back pain undergoing the same experimental task.

Together, our findings establish a neural signature sensitive to both pain exacerbation and pain relief in healthy and clinical samples, providing a potential biomarker for analgesic evaluation and development, and a foundation for future efforts to predict treatment response in pain.

## RESULTS

We analyzed fMRI data from 61 healthy participants (HC-sample), acquired using a previously validated paradigm that enables within-individual modulation of ongoing pain (*12–14*). The ongoing tonic pain state was induced using a capsaicin-based model with individually calibrated heat stimulation. Relative to the tonic pain baseline, pain was systematically modulated in both directions by transient increases (pain exacerbation) and decreases (pain relief) in temperature, alongside a no-change control condition (Fig. 1B). Participants rated pain intensity on a visual analog scale (0–100) for each condition, and subjective pain changes were quantified from condition-wise ratings averaged across trials (Fig. 2A).

**Fig. 1.**
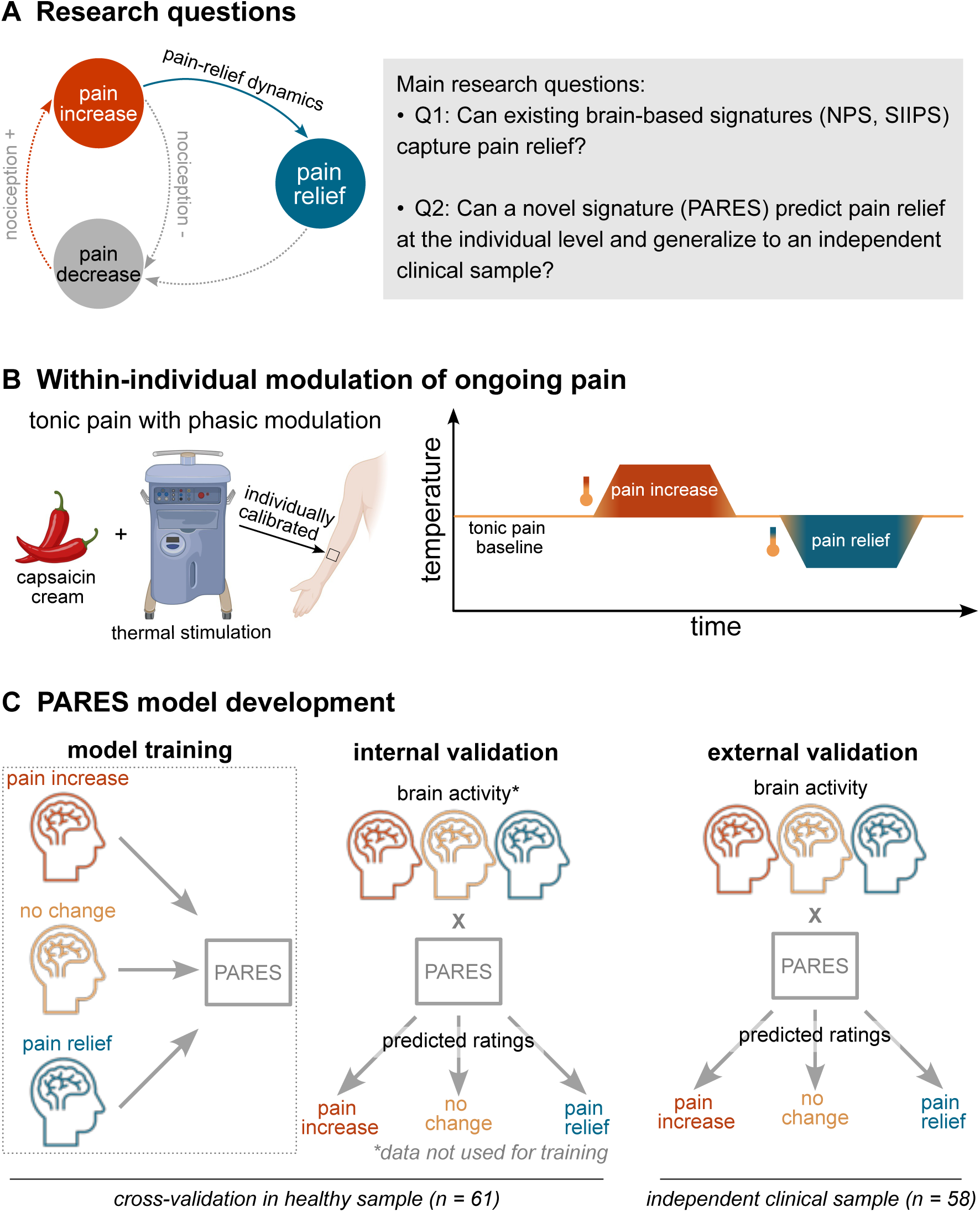
Study rationale, experimental design, and development of PARES. **(A)** Conceptual framework and research questions. **(B)** Within-individual modulation of ongoing pain. Ongoing tonic heat pain was induced using a capsaicin model with individually calibrated thermal stimulation. Relative to the tonic heat pain baseline (“no change”), transient temperature increases and decreases elicited pain exacerbation and pain relief, respectively, allowing within-individual bidirectional pain modulation under the same sensory and experimental context. **(C)** Model development pipeline. PARES was developed on data from 61 healthy participants to predict changes in pain intensity ratings relative to the tonic heat pain baseline. Model performance was evaluated within the healthy sample and externally validated in an independent clinical cohort of n = 58 people with chronic back pain, who underwent the identical fMRI experimental paradigm. NPS, Neurologic Pain Signature; SIIPS, Stimulus Intensity Independent Pain Signature; PARES, PAin and RElief Signature. In (B), the inset “tonic pain with phasic modulation” is partially created in BioRender. https://BioRender.com/3u6jp1r.

**Fig. 2.**
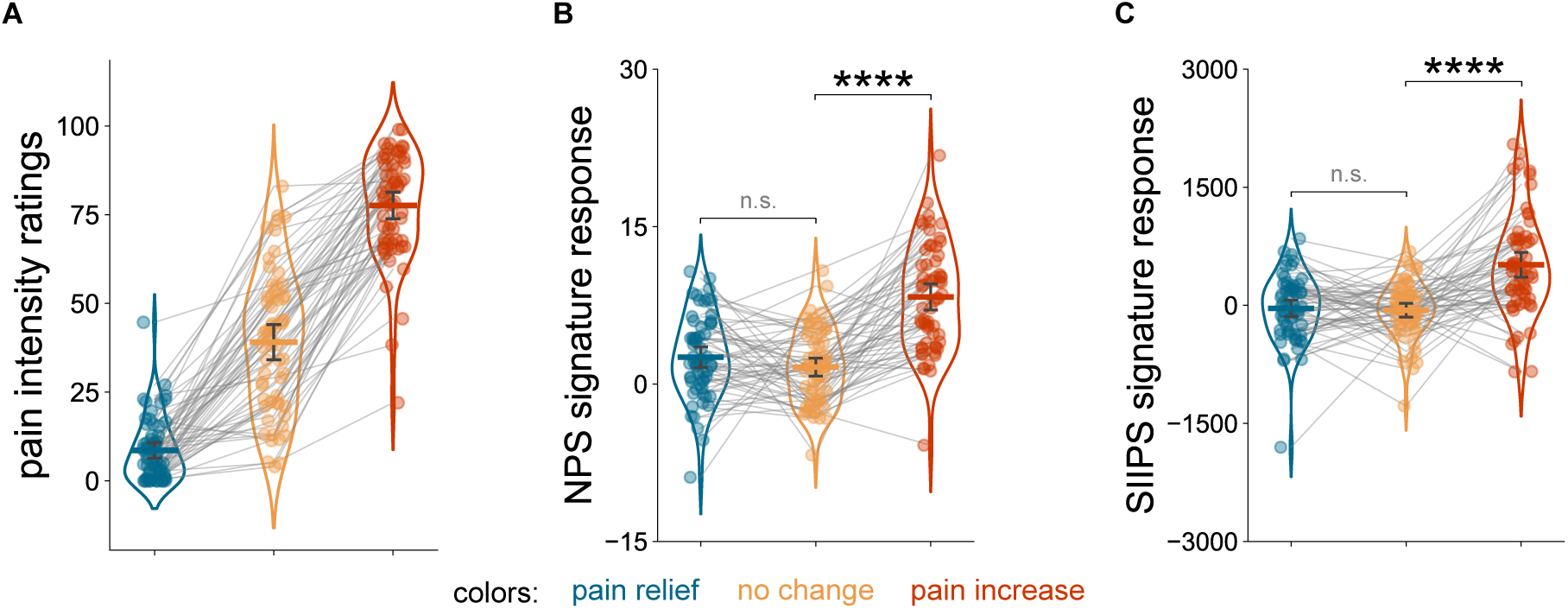
Existing pain signatures detect pain exacerbation but fail to capture pain relief. **(A)** Behavioral pain intensity ratings across experimental conditions in the healthy sample. Relative to the tonic heat pain baseline (“no change”), participants reported lower pain intensity ratings during the pain-relief condition and higher ratings during the pain-increase condition, confirming successful bidirectional modulation of pain. **(B)** Neurologic Pain Signature (NPS) responses across conditions in the healthy sample. The NPS robustly differentiated pain increase from the tonic pain baseline but did not distinguish pain relief from baseline, indicating limited sensitivity to reductions in ongoing pain. **(C)** Stimulus Intensity Independent Pain Signature (SIIPS) responses across conditions in the healthy sample. Similar to the NPS, the SIIPS differentiated pain increase from baseline but failed to detect pain relief. Together, these findings highlight a gap in current brain-based pain models: existing signatures capture pain exacerbation but not the neural representation of decreasing pain. In all panels, each dot represents one participant in one condition, and gray lines connect conditions within participants. Violin plots show value distributions, with horizontal lines indicating means and error bars representing 95% bootstrap confidence intervals. Statistical significance for within-participant contrasts was assessed using paired tests with permutation-based inference. n.s., not significant; ****p < .0001.

### Existing pain signatures do not capture pain relief

We first examined whether the NPS (*5*) and the SIIPS (*8*) capture bidirectional changes in pain perception (Q1, Fig. 1A). Individual brain activation maps were extracted for each condition (see Supplementary Fig. S1) and used to generate condition-wise signature responses. Model performance was evaluated using complementary metrics assessing both between- and within-participant prediction of pain intensity ratings, as well as the ability to differentiate pain increase and pain relief from the tonic pain baseline (see Supplementary Table S1 for details). Statistical significance was assessed using permutation testing (see Methods for details).

We found a statistically significant correlation between pain ratings and NPS predictions, both at the within- and between-participant levels, pooled over conditions (inter-individual: *r* = .422, *p* < .0001; intra-individual: *r* = .555, *p* < .0001). Further, the NPS successfully differentiated the pain increase condition from the “no change” baseline condition at the within-participant level (*t*(60) = 9.483, *p* < .0001), corroborating the NPS’ validity and robustness for capturing within-individual changes in nociceptive activity in response to pain exacerbation. Results are reported in Fig. 2B & Table 1.

**Table 1.**
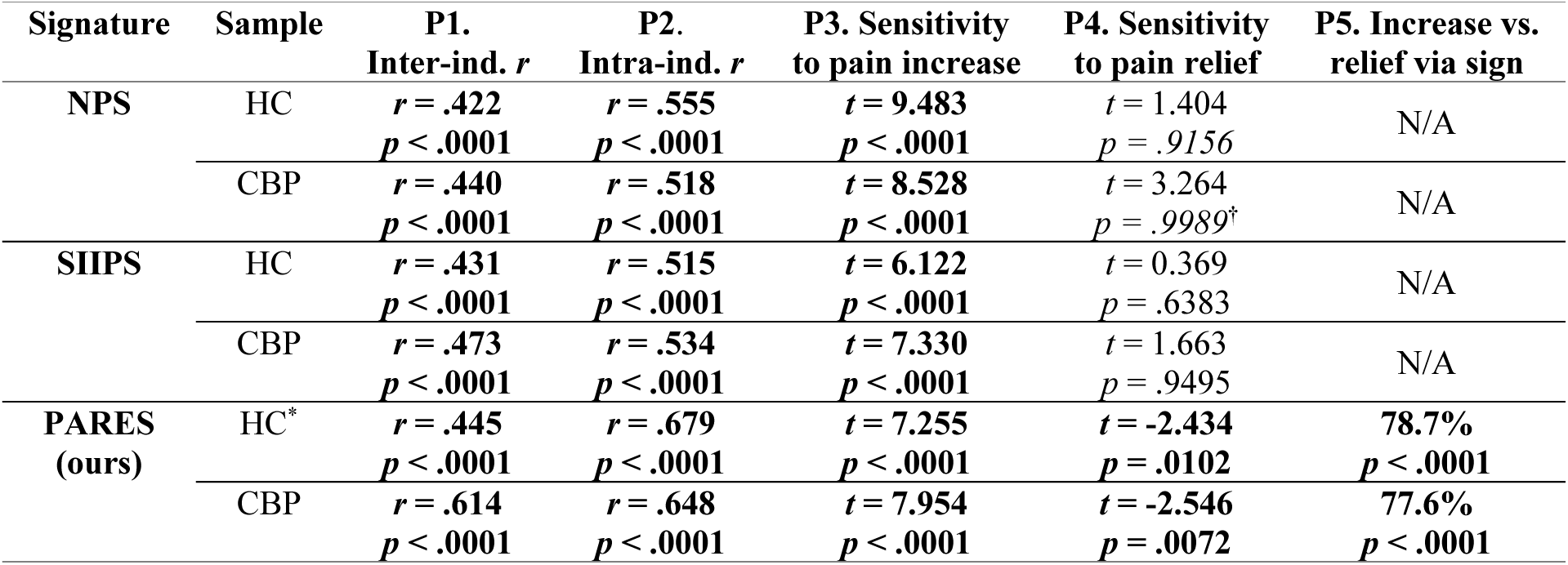
Performance of existing and novel multivariate pain signatures. Performance metrics for the Neurologic Pain Signature (NPS), Stimulus Intensity Independent Pain Signature (SIIPS), and PAin and RElief Signature (PARES). Metrics assessed between-participant prediction (P1), within-participant prediction (P2), sensitivity to pain increase (P3) and pain relief (P4) relative to the tonic pain baseline, and discrimination between pain increase and pain relief based on the sign of the PARES response (P5). For definitions of all performance metrics, see Methods and Supplementary Table S1. Bold values indicate statistical significance under directional (one-tailed) hypothesis testing. Inter-ind. r, between-participant correlation; Intra-ind. r, mean within-participant correlation; HC, healthy controls; CBP, people with chronic back pain; *, internal validation (cross-validated performance measures); ^†^, significance under two-tailed testing in the direction opposite to the hypothesis.

In contrast, the NPS did not distinguish pain relief from the tonic pain baseline under the directional hypothesis of reduced NPS expression during relief (*t*(60) = 1.404, *p* = .9156; paired *t*-test). Instead, exploratory two-sided testing suggested a trend towards the opposite effect direction, with higher rather than lower NPS responses during relief (*p* = .1690; *p* = .0024 in the clinical sample, see Table 1 and Supplementary Fig. S2).

The SIIPS showed a similar pattern. The signature significantly predicted pain ratings across conditions at the between-participant (*r* = .431, *p* < .0001) and the within-participant levels (*r* = .515, *p* < .0001) and differentiated pain increase from the tonic pain baseline (*t*(60) = 6.122, *p* < .0001; paired *t*-test). However, like the NPS, the SIIPS failed to distinguish pain relief from the intra-individual tonic pain baseline (*t*(60) = 0.369, *p* = .6383; Fig. 2C, Table 1).

Together, these findings indicate that existing brain-based pain signatures robustly capture pain exacerbation but fail to detect pain relief.

### Development of a brain-based signature for bidirectional pain prediction

To address this gap, we developed a novel multivariate brain-based signature (PARES) designed to predict bidirectional changes in pain perception (Q2, Fig. 1A). Rather than estimating pain intensity along a single direction, PARES was trained to predict changes in pain relative to the tonic pain baseline (see Methods and Fig. 1C). This change-based formulation allowed a single model to capture both pain increase and pain relief within a unified framework, with positive predictions indicating pain increase and negative predictions indicating pain relief.

Model performance was evaluated internally (within the training dataset) using cross-validation and tested for generalization in an independent cohort of people with chronic back pain (CBP-sample, n = 58). Detailed model specifications, validation procedures, and additional analyses are provided in the Methods and Supplementary Methods. Performance of the NPS and SIIPS in the CBP-sample is shown in Table 1 and Supplementary Fig. S2.

### PARES predicts both pain increase and relief

During internal performance estimation (cross-validated HC-sample), our model predicted differential pain intensity ratings with a large effect size, at both the between-participant (*r* = .445, *p* < .0001, explained variance: R^2^ = 19.8%) and within-participant levels (*r* = .679, *p* < .0001, explained variance: R^2^ = 46.1%; Fig. 3A, Table 1).

**Fig. 3.**
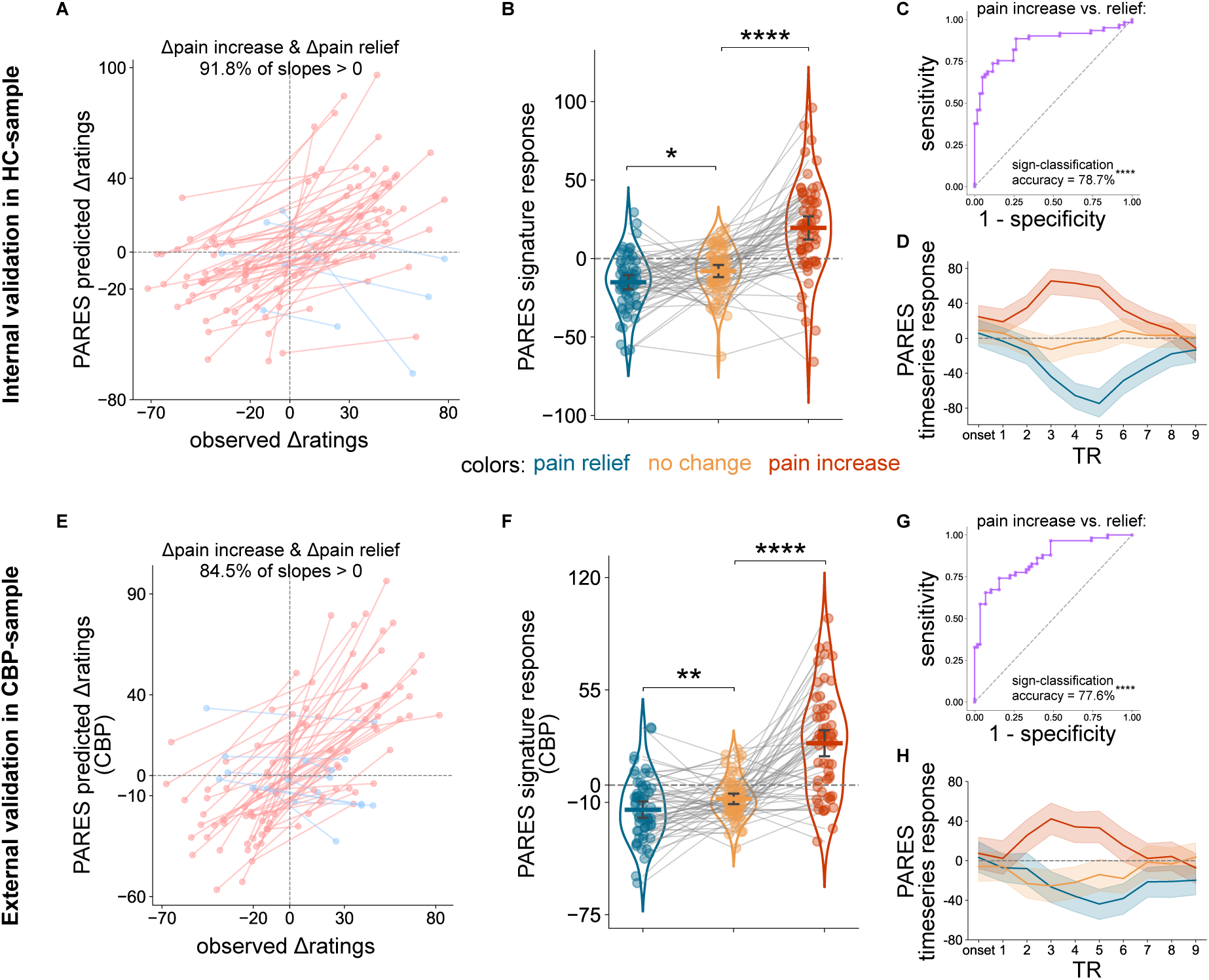
PARES robustly captures both pain exacerbation and pain relief. **(A)** Internal validation in the healthy participants (HC-sample). Cross-validated PARES predictions tracked within-participant pain intensity rating changes relative to the no-change condition, showing that the model captured both pain exacerbation and pain relief on a common continuous scale. Most participant-level slopes were positive (pink color), indicating that higher PARES responses were generally associated with greater pain increase and lower responses with greater relief. **(B)** Condition-wise PARES responses in the HC-sample. Relative to the no-change condition, PARES responses were significantly elevated during pain increase and reduced during pain relief. **(C)** Receiver operating characteristic (ROC) analysis in the HC-sample. The sign of the PARES response alone distinguished pain increase from pain relief with high accuracy, consistent with a calibrated representation centered around zero. **(D)** Time-course expression of PARES in the HC-sample. PARES responses diverged following stimulus onset, with sustained positive expression during pain increase and negative expression during pain relief (methods see Supplementary Methods 1.4). **(E)** External validation in the chronic back pain (CBP) sample. Applied without retraining, PARES again tracked observed pain rating changes in an independent clinical cohort, demonstrating generalizability beyond the healthy training sample. **(F)** Condition-wise PARES responses in the CBP-sample. As in the HC-sample, PARES responses were significantly elevated during pain increase and reduced during pain relief relative to the no-change condition. **(G)** ROC analysis in the CBP-sample. Sign-based classification successfully distinguished the two states. **(H)** Time-course expression of PARES in the CBP-sample. Patients showed a temporal response profile similar to healthy participants, with positive expression during pain increase and negative expression during pain relief. In (A) and (E), each point represents one condition from one participant; line segments connect pain-increase and pain-relief observations within participants; pink indicates positive within-participant slopes and blue negative slopes. In (B) and (F), each dot represents one participant and gray lines connect conditions within participants; violin plots show distributions, with the central horizontal line indicating means and error bars representing 95% bootstrap confidence intervals. In (D) and (H), solid lines indicate condition-wise mean PARES time-course responses and shaded bands indicate 95% confidence intervals. TR, repetition time; *p < .05; **p < .01; ****p < .0001.

PARES further differentiated bidirectional changes in pain perception across conditions. Importantly, because the model was trained to predict continuous pain intensity changes without being provided with categorical condition labels, these findings suggest that PARES captured pain increase and pain relief within a common representational space. Relative to the tonic pain baseline, condition-wise cross-validated PARES responses were significantly higher during pain increase (*t*(60) = 7.255, *p* < .0001; paired *t*-test) and significantly lower during pain relief (*t*(60) = -2.434, *p* = .0102; paired *t*-test) (Fig. 3B). The sign of the PARES-score alone was sufficient to distinguish pain increase from pain relief, with 78.7% classification accuracy pooled across participants (*p* < .0001), 75.4% sensitivity and 82.0% specificity. The receiver operating characteristic (ROC) analysis yielded an area under the curve (AUC) of 86.0% (Fig. 3C).

### PARES generalizes to an independent clinical sample

External validation in an independent cohort of individuals with chronic back pain, who underwent the same experimental paradigm of transient pain increase and decrease, corroborated the internal cross-validated performance estimates of PARES. When applied to the CBP-sample, the model significantly predicted differential pain ratings at both the between-participant (*r* = .614, *p* < .0001, explained variance: R^2^ = 37.7%) and the within-participant levels (*r* = .648, *p* < .0001, explained variance: R^2^ = 42.0%; Fig. 3E, Table 1). PARES further differentiated bidirectional changes in pain perception in the clinical sample. Relative to the tonic baseline, PARES responses were significantly higher during pain increase (*t*(57) = 7.954, *p* < .0001; paired *t*-test) and significantly lower during pain relief (*t*(57) = -2.546, *p* = .0072; paired *t*-test) (Fig. 3F).

The sign of the PARES response further yielded significant classification accuracy for distinguishing pain increase from pain relief (77.6%, *p* < .0001), with 74.1% sensitivity, 81.0% specificity and AUC of 85.8% (Fig. 3G). These results suggest that PARES predictions generalize to independent clinical data, preserving the interpretation of positive responses as pain increase, negative responses as pain relief and values near zero as minimal change in pain perception.

### Distinct and shared predictors of pain increase and pain relief

To examine whether PARES relies on shared or distinct neural predictors for pain increase and relief, we characterized the condition-specific contribution of brain regions that significantly contributed to model prediction (false discovery rate, FDR *q* < 0.05). For details, see Methods and Supplementary Fig. S3.

Our analysis revealed partially dissociable neural patterns for pain increase and relief (Table 2). Pain exacerbation was predominantly associated with increased activity in regions linked to nociceptive and somatosensory processing, including the secondary somatosensory cortex (S2), mid/posterior insula (m/pINS), thalamus, dorsal anterior cingulate cortex (dACC), supplementary motor area, precentral cortex, cerebellum and brainstem. In contrast, pain relief more strongly engaged frontoparietal and valuation-related regions, including the orbitofrontal cortex (OFC), intraparietal sulcus (IPS), inferior parietal lobule (IPL), ventromedial prefrontal cortex (vmPFC), and parts of lateral parietal cortex, while some default mode regions such as the posterior cingulate cortex showed deactivation during relief (Fig. 4A-B, Fig. 5). Interestingly, the periaqueductal gray (PAG) contributed the strongest positive influence to model prediction (i.e., the highest peak-level multivariate weight) and showed opposite response profiles across conditions, with increased activity during pain exacerbation (Haufe-*t* = 2.528) and decreased activity during relief (Haufe-*t* = -1.516; Fig. 4C), consistent with its role in pain facilitation and descending control (*15, 16*). Detailed results see Supplementary Table S2-3.

**Fig. 4.**
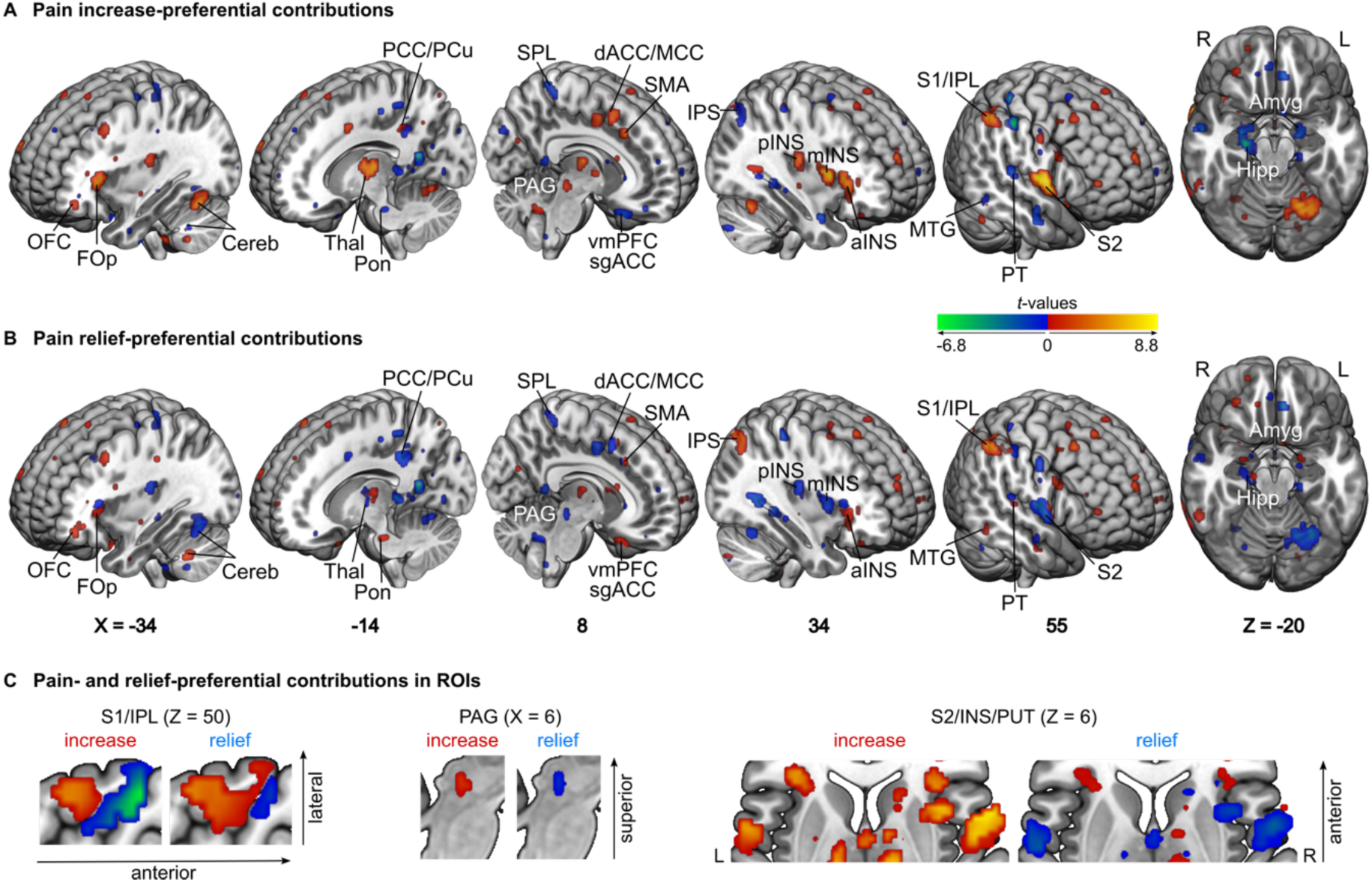
PARES combines predictors with preferential contributions to pain increase and pain relief. **(A)** Pain increase-preferential contributions to PARES predictions. Restricting the analysis to voxels contributing significantly to the multivariate model (false discovery rate, FDR q < .05) and applying a Haufe transformation for pain increase versus no change revealed prominent expression in nociception-associated regions, including the S2, mid/posterior insula, thalamus, dACC/MCC, SMA, and brainstem/PAG. This pattern is consistent with enhanced nociceptive processing during transient pain exacerbation. **(B)** Pain relief-preferential contributions to PARES predictions. In contrast, pain relief showed a partially distinct pattern, with relatively stronger involvement of frontoparietal and valuation-related regions, including the OFC, IPS, vmPFC, and lateral parietal cortex, alongside reduced expression in parts of the nociceptive system. Together, these maps indicate that PARES captures partly dissociable neural contributions to worsening and decreasing pain. **(C)** Regions-of-interest (ROIs) zooms highlighting representative local dissociations within the PARES pattern. Insets illustrate heterogeneous contributions in the combined S1/IPL cluster, the PAG, and the S2/insula/putamen region, emphasizing that neighboring voxels can show different preferential expression for pain increase and pain relief. Warm colors indicate positive Haufe t-values and cool colors indicate negative Haufe t-values for the indicated contrasts. The same t-value range is used for the pain-increase and pain-relief maps to facilitate direct visual comparison. Labels indicate representative anatomical locations. OFC, orbitofrontal cortex; FOp, frontal operculum; Cereb, cerebellum; Thal, thalamus; Pon, pons; PCC, posterior cingulate cortex; PCu, precuneus; PAG, periaqueductal gray; vmPFC, ventromedial prefrontal cortex; sgACC, subgenual anterior cingulate cortex; SPL, superior parietal lobule; dACC, dorsal anterior cingulate cortex; MCC, midcingulate cortex; SMA, supplementary motor area; IPS, intraparietal sulcus; pINS, posterior insula; mINS, mid-insula; aINS, anterior insula; MTG, middle temporal gyrus; PT, planum temporale; S2, secondary somatosensory cortex; S1, primary somatosensory cortex; IPL, inferior parietal lobule; Amyg, amygdala; Hipp, hippocampus; PUT, putamen; L, left; R, right.

**Fig. 5.**
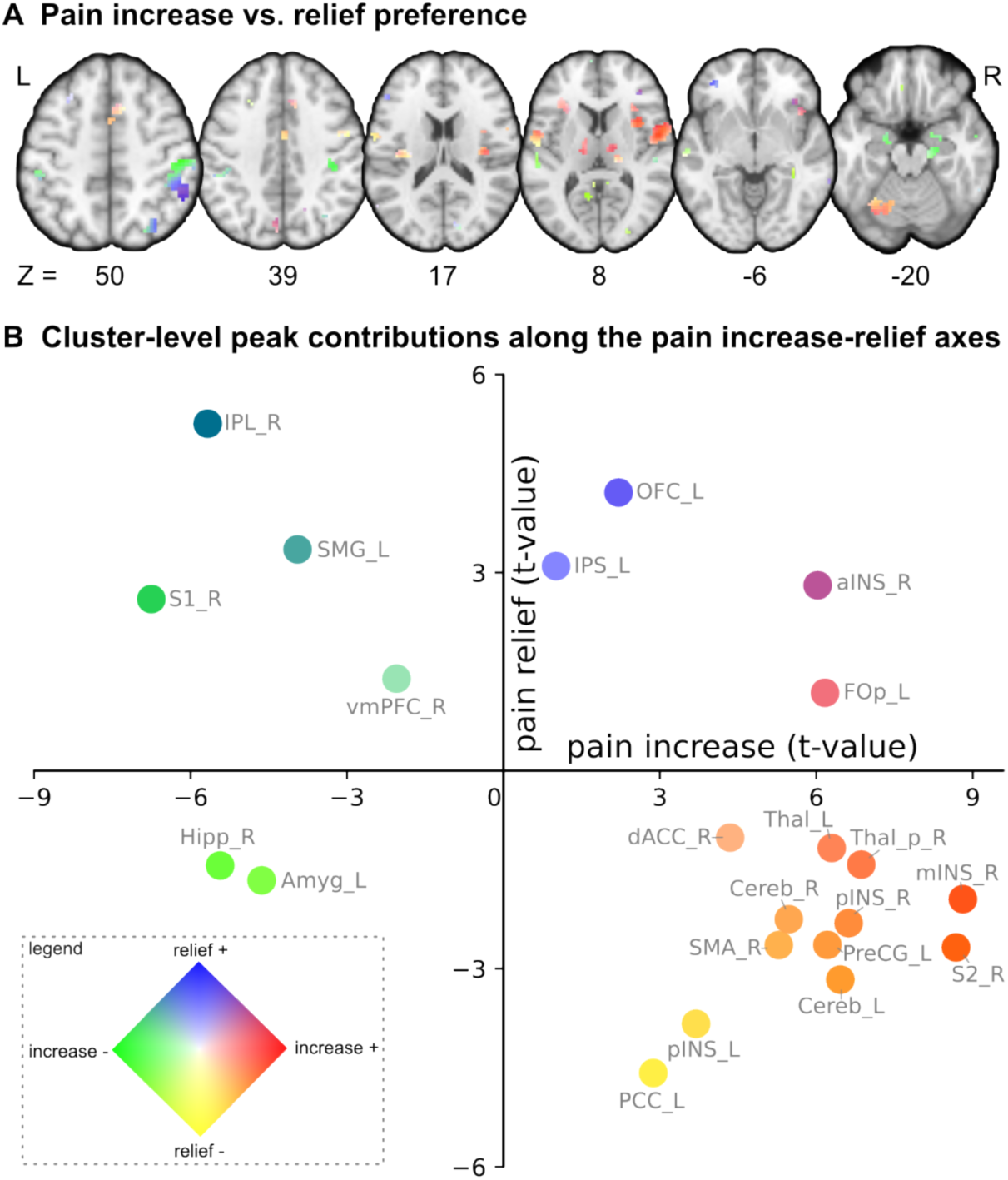
Two-dimensional visualization of preferential contributions to pain increase and pain relief within the PARES pattern. **(A)** Voxel-level pain increase and relief preferential Haufe-weights displayed in a two-dimensional color space. Each voxel is colored according to its pair of condition-specific Haufe t-values, as encoded in the two-dimensional colormap on panel (B), with the horizontal dimension reflecting expression during pain increase versus no change and the vertical dimension reflecting expression during pain relief versus no change. Voxels contributing to PARES spanned multiple response profiles, including preferential expression during pain increase (green or red), preferentially expressed during relief (blue or yellow), and shared directional responses across both conditions with different magnitudes (purple: both positive, orange: positive in increase negative in relief). **(B)** Cluster-level summary of preferential contributions within the PARES pattern. For each FDR-significant cluster, the signed maximal absolute Haufe t-value for pain increase is plotted on the x-axis and the corresponding value for pain relief on the y-axis. Regions more strongly associated with pain increase were concentrated in nociception-associated areas, including the S2, mid/posterior insula, thalamus, dACC, SMA, precentral cortex, cerebellum, and PCC. In contrast, regions more strongly associated with relief included frontoparietal and valuation-related areas such as the OFC, IPS, IPL, vmPFC, and SMG. Some regions showed similar directional responses across both conditions but with different magnitudes, indicating overlapping but non-identical neural contributions to pain increase and pain relief. Note that a large cluster with heterogeneous pain increase-relief contributions extended across S1 and IPL regions (Fig. 4C first slice). For the scatter plot, the cluster was manually subdivided into the S1and IPL regions based on the Juelich histological brain atlas (17) (see Methods and Supplementary Fig. S5 for details). In (A), only voxels with significant multivariate contributions to PARES are shown (bootstrap-derived FDR q < .05). In (B), each point represents one cluster, labeled by anatomical region and hemisphere. Colors in both panels correspond to the two-dimensional colormap shown in the legend: red, activation during pain increase; green, deactivation during pain increase; blue, activation during pain relief; yellow, deactivation during pain relief; intermediate colors indicate combinations of these response directions. OFC, orbitofrontal cortex; IPS, intraparietal sulcus; aINS, anterior insula; FOp, frontal operculum; IPL, inferior parietal lobule; S1, primary somatosensory cortex; SMG, supramarginal gyrus; vmPFC, ventromedial prefrontal cortex; Amyg, amygdala; Hipp, hippocampus; dACC, dorsal anterior cingulate cortex; Thal_p, posterior thalamus; mINS, mid-insula; pINS, posterior insula; Cereb, cerebellum; SMA, supplementary motor area; S2, secondary somatosensory cortex; PreCG, precentral gyrus; PCC, posterior cingulate cortex; L, left; R, right.

**Table 2.** Clusters with predictive contributions surviving FDR correction (FDR q < .05; KE > 10 voxels). abs., absolute; Thal_p, posterior thalamus; S2, secondary somatosensory cortex; pINS, posterior insula; PreCG, precentral gyrus; Cereb, cerebellum; mINS, mid-insula; aINS, anterior insula; SMA, supplementary motor area; dACC, dorsal anterior cingulate cortex; PCC, posterior cingulate cortex; FOp, frontal operculum; M1, primary motor cortex; MFG, middle frontal gyrus; PCu, precuneus; vmPFC, ventromedial prefrontal cortex; Hipp, hippocampus; LG, lingual gyrus; OFC, orbitofrontal cortex; PT, planum temporale; SMG, supramarginal gyrus; Amyg, amygdala; sgACC, subgenual anterior cingulate cortex; S1, primary somatosensory cortex; IPS, intraparietal sulcus; SFG, superior frontal gyrus; IPL, inferior parietal lobule; MTG, middle temporal gyrus; V2, secondary visual cortex.

| Cluster size (mm<sup>3</sup>) | X | Y | Z | Region | Multivariate weight (peak) | FDR $q$ (peak) | increase | relief |
| --- | --- | --- | --- | --- | --- | --- | --- | --- |
| | | | | | | | Signed max abs. Haufe-$t$ | Signed max abs. Haufe-$t$ |
| positive predictive weights |  |  |  |  |  |  |  |  |
| 954 | 18 | -23 | 8 | Thal_p | 0.0176 | < .0001 | 6.862 | -1.424 |
| 2931 | 55 | 2 | 11 | S2 | 0.0171 | .0001 | 8.679 | -2.679 |
| 659 | 35 | -16 | 17 | pINS | 0.0171 | < .0001 | 6.620 | -2.310 |
| 676 | -37 | -18 | 17 | pINS | 0.0165 | < .0001 | 3.693 | -3.836 |
| 1960 | -59 | -3 | 14 | PreCG | 0.0162 | < .0001 | 6.213 | -2.640 |
| 1890 | -29 | -63 | -17 | Cereb | 0.0152 | < .0001 | 6.460 | -3.167 |
| 659 | -14 | -8 | 11 | Thal | 0.0147 | .0004 | 6.295 | -1.173 |
| 971 | 33 | 9 | 8 | mINS | 0.0147 | < .0001 | 8.810 | -1.947 |
| 1248 | 30 | 24 | 8 | aINS | 0.0145 | .0073 | 6.027 | 2.807 |
| 225 | 13 | -43 | -51 | brainstem | 0.0131 | < .0001 | 2.320 | -3.220 |
| 1214 | 8 | 14 | 50 | SMA | 0.0125 | .0044 | 5.281 | -2.643 |
| 190 | 11 | 24 | 39 | dACC | 0.0121 | .0188 | 4.349 | -1.015 |
| 294 | -4 | -48 | 3 | PCC | 0.0118 | .0004 | 2.875 | -4.582 |
| 728 | -29 | 24 | 11 | FOp | 0.0116 | .0011 | 6.166 | 1.179 |
| 329 | 30 | -58 | -23 | Cereb | 0.0114 | < .0001 | 5.473 | -2.252 |
| 572 | 48 | -3 | 42 | PreCG | 0.0114 | < .0001 | 4.574 | -2.205 |
| 208 | 35 | -30 | 56 | M1 | 0.0112 | < .0001 | -2.015 | -2.660 |
| 797 | -27 | 24 | 39 | MFG | 0.0099 | .0241 | 2.543 | 1.998 |
| 225 | -4 | -73 | 39 | PCu | 0.0088 | .0152 | 5.401 | 2.159 |
| negative predictive weights |  |  |  |  |  |  |  |  |
| 485 | 8 | 24 | -28 | vmPFC | -0.0176 | < .0001 | -2.050 | 1.393 |
| 1127 | 18 | -20 | -20 | Hipp | -0.0155 | .0002 | -5.433 | -1.438 |
| 208 | 16 | -75 | 0 | LG | -0.0131 | .0003 | -1.966 | -1.359 |
| 242 | -34 | 44 | -6 | OFC | -0.0125 | .0214 | 2.209 | 4.212 |
| 277 | 53 | -28 | 11 | PT | -0.0122 | .0140 | -2.496 | -1.043 |
| 988 | -44 | -33 | 39 | SMG | -0.0119 | .0112 | -3.947 | 3.350 |
| 294 | -19 | -6 | -23 | Amyg | -0.0118 | < .0001 | -4.637 | -1.663 |
| 208 | 13 | 32 | -9 | sgACC | -0.0117 | .0060 | -2.418 | 1.690 |
| 646 | 60 | -23 | 50 | S1 | -0.0114 | .0004 | -6.752 | 2.599 |
| 416 | -29 | -25 | 67 | M1 | -0.0111 | .0054 | -4.254 | -2.428 |
| 208 | -39 | -48 | 33 | IPS | -0.0107 | .0001 | 1.009 | 3.098 |
| 346 | 23 | 37 | 28 | SFG | -0.0106 | .0004 | 1.318 | 2.419 |
| 1343 | 60 | -25 | 50 | IPL | -0.0105 | .0004 | -5.671 | 5.256 |
| 208 | -52 | -23 | 33 | S1 | -0.0105 | .0082 | -1.806 | 1.645 |
| 693 | 65 | -50 | -9 | MTG | -0.0104 | .0456 | 2.180 | 2.503 |
| 294 | -54 | -28 | 8 | PT | -0.0102 | .0161 | -3.483 | -2.541 |
| 468 | 13 | -83 | 19 | V2 | -0.0100 | .0175 | -1.712 | 1.183 |
| 641 | 33 | -68 | 42 | IPS | -0.0093 | .0083 | -2.422 | 3.657 |

Meta-analytic decoding (Neurosynth) associated our pain increase Haufe-weight map with terms related to nociception and somatosensation, whereas the pain relief Haufe-weight map was preferentially associated with cognitive control, attention, working memory, and uncertainty (Supplementary Fig. S4).

These results suggest that pain increase and relief are not represented as opposite ends of a single nociceptive axis. Instead, PARES integrates overlapping but partially distinct neural contributions to capture bidirectional changes in pain perception.

## DISCUSSION

Using a tonic pain model that enabled controlled induction of both pain exacerbation and relief, we found that (i) established brain-based pain signatures, the NPS and the SIIPS, detect pain increases but fail to capture the subjective experience of pain relief; (ii) our newly developed multivariate signature PARES reliably predicts bidirectional changes in pain perception and generalizes to an independent cohort of people with chronic back pain; and (iii) pain increase and relief involve overlapping but partially distinct neural contributions.

The inability of both the NPS and the SIIPS to capture pain relief highlights an important limitation of current brain-based models of pain. Whilst both signatures have been extensively validated for tracking pain intensity (*5, 6, 8, 18–20*), our findings suggest that reductions in ongoing pain engage additional neural processes that are not sufficiently represented in existing models. This interpretation is consistent with prior evidence showing limited sensitivity of the NPS to contextual and modulatory influences on pain, including placebo analgesia, cognitive regulation and social context (*5, 7, 8, 21–23*). Importantly, failure of the SIIPS to detect relief further suggests that the missing component is not merely higher-order modulation per se, but the representation of relief as a distinct neurobiological state.

PARES was developed to model these bidirectional changes in pain perception. The signature reliably captured both pain exacerbation and pain relief across both healthy participants and people with chronic back pain. This distinction to previous brain-based signatures is of high clinical relevance because it positions PARES as a potential tool for monitoring treatment efficacy and identifying impaired relief-related pain regulation. By capturing bidirectional changes in pain perception, PARES may complement existing nociceptive pain biomarkers and provide a more clinically relevant measure of treatment-related improvement.

The regional contributions to PARES further support the idea that pain relief engages processes beyond reduced nociceptive signaling (Fig. 5): While pain increase was more strongly associated with canonical nociceptive and somatosensory regions including the thalamus, dACC, mid and dorsal-posterior insula commonly implicated in pain processing (*24, 25*), pain relief more strongly involved valuation-related regions consistent with prior work on higher-order cognitive control, attention allocation and affective valuation in pain (*26–29*). Because PARES integrates both nociception-related and relief-related representations within a single predictive framework, it is able to capture bidirectional changes in pain perception.

Particularly noteworthy was the contribution of the OFC and vmPFC to the relief-related component of PARES. Both regions have repeatedly been implicated in valuation, expectancy-related processing, and adaptive updating of motivational significance (*30–33*). This interpretation aligns with reinforcement-learning and opponent-process accounts proposing that relief from pain is represented as an appetitive motivational state rather than merely the absence of nociceptive input (*11, 12*). It is further supported by evidence that tonic pain suppresses medial OFC activity associated with reward-related valuation processes (*34, 35*), suggesting that relative pain relief may engage neural systems involved in affective and motivational re-evaluation. Together, these findings support the view that relief-related representations within PARES reflect processes such as evaluative updating, attentional reorientation, expectancy-related processing, and endogenous regulation as pain decreases.

The dissociation between pain increase and relief at the neural level may also have important implications for understanding chronic pain. Chronic pain has increasingly been linked to not only amplified nociceptive processing, but also to alterations in endogenous regulatory, motivational, and affective systems involved in pain modulation (*36, 37*). By explicitly modeling relief-related representations, PARES may provide a framework for investigating how such processes contribute to the persistence or resolution of pain. In particular, altered neural responses during pain relief may reflect disruptions in evaluative, motivational, or regulatory mechanisms that normally support adaptive updating as pain decreases. For example, reduced PARES responses during expected relief periods could indicate disrupted descending modulation or reward-based pain regulation – processes thought to contribute to chronic pain maintenance (*2, 10*).

Our results further demonstrate that pain relief can be robustly captured within an experimentally controlled yet clinically relevant framework. Building on this foundation, future studies should evaluate the extent to which PARES generalizes beyond experimentally induced pain modulation to spontaneous fluctuations in clinical pain and less constrained neuroimaging paradigms. Emerging approaches that capture spontaneous pain dynamics may provide an important bridge between controlled experimental settings and real-world pain experiences (*38, 39*). Such efforts will be critical for translating experimentally derived neural signatures into tools that can inform the assessment and mechanistic understanding of pain relief in clinical settings.

Beyond its translational potential as a monitoring biomarker (*40–42*), PARES provides a framework for identifying neural processes associated with pain relief. The preferential involvement of valuation- and cognitive control-related systems highlights these processes as promising targets for future mechanistic and interventional research aimed at enhancing adaptive pain regulation and improving treatment outcomes.

In conclusion, our findings indicate that pain increase and pain relief are represented by partially dissociable brain patterns, with pain increase weighted toward regions associated with nociceptive processing and relief more strongly engaging frontoparietal and valuation-related representations. By capturing bidirectional changes in pain perception within an ongoing tonic pain model, PARES extends brain-based pain biomarkers to a more clinically relevant and ecologically valid representation of pain dynamics. PARES may provide a mechanistically informed readout of target engagement in analgesic development, facilitate treatment monitoring and stratification in chronic pain, and highlight neural processes that may guide the development of future pharmacological and non-pharmacological interventions.

## MATERIALS AND METHODS

### Participants

The study involved a total of 119 participants including n = 61 healthy individuals (HC-sample) and n = 58 individuals with chronic back pain (CBP) (CBP-sample). Data from the HC-sample served as a cross-validated dataset for the PARES model development, and the CBP-sample was used for external validation. Participants were screened for eligibility by trained experimenters, using screening forms for MRI and safety checks (e.g., MRI contraindications, pregnancy or breastfeeding, capsaicin allergy; for details see Supplementary Methods 1.1). Further on-site screening of eligible patients was performed by physicians specialized in pain medicine through medical history and/or clinical examinations. Patients experiencing non-specific CBP were included. The study was registered at the German register for clinical trials and approved by the local Ethics Committee (16-7248-BO) of the Medical Faculty, University Duisburg-Essen. Participants gave written informed consent before the start of the experiment and could withdraw from the study at any time. Details can be found in Supplementary Methods 1.1 and (*14*).

### Procedure and experimental paradigm

Both samples underwent the same experimental procedure. Detailed descriptions of the full procedure, paradigm and temperature calibration are provided in Supplementary Methods 1.2, 1.3 and (*14*).

On the first day, individual temperature levels for heat stimulations were determined using a validated capsaicin-induced tonic heat pain model (*13*) corresponding to subjective pain intensity ratings on a visual analog scale (VAS). On the second day, following individual re-calibration, participants underwent fMRI scanning while receiving continuous tonic heat stimulation with transient temperature changes representing pain increase, pain relief, or no change conditions. The order of condition was pseudorandomized and varied across participants. During the experimental task, visual geometric cues preceded each temperature event and served as a cue indicating the upcoming condition (i.e., pain increase, pain relief, no change). Pain intensity was rated in response to the question “How painful was this temperature stimulus?” using VAS 0-100 (anchors: 0 = “not painful at all”, 100 = “unbearably painful”). Ratings were collected following every fourth stimulus of the same type, yielding three ratings per condition. We averaged the individual-level pain intensity ratings across the three timepoints for each condition (Fig. 2A, Supplementary Fig. S2A).

After the functional runs, high-resolution structural (T1-weighted) images were acquired.

### Preprocessing and GLM analysis of fMRI data

Neuroimaging data were preprocessed using fMRIPrep v22.1.1 (*43*). Preprocessing of participants’ T1-weighted images included skull-stripping, spatial normalization and brain tissue segmentation. Functional images were preprocessed by co-registration with individual T1-weighted reference image, motion correction, slice-timing correction and normalization to MNI152NLin2009cAsym standard space. Physiological regressors were calculated using component-based noise correction (CompCor), and the first 6 anatomical components (aCompCor) for each participant were added as nuisance regressors to the individual design matrix in the first-level analysis (*44*). Functional images were spatially smoothed using a Gaussian kernel with 6mm full-width at half-maximum (FWHM).

Univariate general linear model (GLM) analyses were conducted using SPM12 (https://www.fil.ion.ucl.ac.uk/spm/). The individual design matrices included regressors-of-interest modelling pain stimulation conditions, i.e., pain increase, relief and no change (duration: 8s), additional regressors coding for the anticipation phase during cue presentations, the rating phase and the 6 CompCor regressors described above.

### Multivariate signature evaluation

We evaluated two established multivariate pain signatures, the NPS and the SIIPS in both datasets by calculating the dot-product of model-weights with individual-level brain activation beta-maps for each condition (pain increase, pain relief, and no change), yielding one predicted score per individual per condition.

Predictive performance of the signatures was assessed using four primary approaches (Supplementary Table S1). Pooled inter-individual performance (P1): we quantified the models’ overall ability to capture pain regardless of conditions between participants by computing Pearson correlations between predicted and observed ratings. Pooled intra-individual performance (P2): we also assessed how well the models capture within-individual variations in ratings by averaging within-individual observation-prediction correlations across participants. Sensitivity to pain increase and pain relief (P3, P4): we evaluated each model’s ability to differentiate pain increase and pain relief from the no change condition within participants by paired *t*-tests.

Statistical significance for all performance metrics was determined via permutation testing with 10,000 iterations with one-tailed tests, e.g., whether the NPS response increased during pain increase and decreased during pain relief relative to no change. We exploratorily performed two-tailed tests to assess whether the insignificance brings a “paradoxical” effect, i.e., the opposite direction than expected.

### Development of the PARES

Using brain activation beta-maps during the three conditions from 61 healthy individuals, we developed the PARES that predicted changes in subjective pain intensity ratings associated with pain increase and relief, relative to the individual-level tonic heat pain baseline (Δpain increase, Δpain relief) (Fig. 1C). The model was also trained on activation patterns from the no-change condition, for which the corresponding prediction target was assigned as zero for each participant. During training, the model was not provided with information about which condition each beta-map originated from. However, since the Δratings were computed as differences from individual tonic pain baseline, in case of successful model training, we expected that the sign of the predictions will implicitly capture the underlying affective direction, e.g., positive values reflect pain increase, and negative values reflect pain relief. This approach allowed the PARES to learn continuous representations of pain increase and relief dynamics without explicit categorical supervision and, at the same time, mitigated the effect of epiphenomena potentially accompanying all stimulation blocks but independent of condition (e.g., shared saliency effects).

We implemented the machine learning framework using a Ridge regression model (L2-regularized linear regression), optimized to minimize overfitting given high dimensionality of fMRI data. Model training and evaluation were conducted via a nested leave-one-participant-out cross-validation scheme to ensure unbiased performance estimation and to avoid information leakage between training and testing sets. In each outer loop, data from one participant were held out for testing, while the remaining participants were used for model training. Within the training set, an inner cross-validation loop was performed to identify the optimal regularization from a predefined hyperparameter grid (alpha: 0.05-5.0). The model with the best-fitting hyperparameter was then applied to the left-out participant in the outer loop.

### Evaluating the predictive performance of the PARES

We evaluated the predictive performance of the PARES with the same measures we used for the NPS and SIIPS evaluation (P1-P4). Furthermore, as the PARES yields signed scores relative to the no change condition, we tested whether the sole sign of the predicted value could distinguish pain increase from pain relief (P5) by pooling predictions across both conditions and all participants, i.e., positive scores were treated as predicting pain increase and negative scores as predicting relief. Classification performance for pain increase vs. relief was quantified by sign-based metrics (accuracy, sensitivity, specificity) and by ROC curves and AUC values.

### Identifying significant PARES predictors

To assess the significance of predictive weights, we performed bootstrap tests with 10,000 iterations. In each iteration, participants were randomly resampled with replacement, ensuring that all three conditions for a given participant were sampled together. For each bootstrap sample, the beta-maps and corresponding Δratings were extracted. Ridge regression model (alpha = 5.0, given the best hyperparameter from model training) was fit to predict ratings from the beta-maps. The resulting predictive weights (model coefficients) were stored.

To evaluate significance of voxel-level bootstrap distributions, we estimated empirical one-tailed *p*-values. However, as bootstrap distributions may be heavy-tailed (i.e., the minimum *p*-values are clamped by the number of bootstrap iterations), applying false discovery rate (FDR) correction on these bootstrapped *p*-values may be conservative (*45*). Thus, we fitted a generalized pareto distribution (GPD) (*46*) to the distribution tails in every voxel close to FDR-significance (uncorrected *p*_one-tailed_ < .001, separately for positive and negative weights). The optimal tail ratio was selected from a pre-defined set of values (from 0.5% to 5% of the bootstrap distribution, with increment of 0.45%), based on goodness of fit, as measured by the Kolmogorov-Smirnov statistic (*47*). *P*-values were then approximated based on the fitted Pareto distribution:

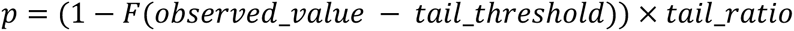

Where F(x) is the Cumulative Distribution Function of the generalized Pareto distribution.

We applied FDR correction on the resulting GPD-based *p*-values, and voxels that survived FDR correction (FDR *q* < .05) were considered significant in the PARES predictive pattern. We visualized the significant voxels for positive and negative weights separately, indicating positive and negative associations between brain activity and Δratings, respectively (Supplementary Fig. S3).

As a supplementary analysis, we also performed a region-level feature importance analysis, which yielded results highly coherent with the voxel-wise analysis (for details see Supplementary Methods 1.6, Supplementary Results 2.4 and Supplementary Fig. S6).

### Disentangling pain increase- and relief-preferential regions

Beyond identifying voxels that significantly contributed to the predictive model, to facilitate signature interpretability (*48*), we further characterized preferential contributions to pain increase and relief using condition-specific Haufe transformations (*49*). For each significant voxel (FDR *q* < .05) in the PARES multivariate pattern, we fitted a linear mixed-effects model to the condition-wise beta-values, with condition (pain increase/relief vs. no change) as a fixed effect and subject as a random intercept. This yielded voxel-wise Haufe *t*-values quantifying univariate BOLD activity differences between conditions (both conditions relative to no-change). No additional statistical threshold was applied, as the voxels investigated were already restricted to those surviving FDR correction.

We visualized regional Haufe-weights in three complementary approaches. First, we displayed voxel-wise Haufe *t*-maps for pain increase and relief contrasts separately and showed on matched brain slices to facilitate comparison between conditions (Fig. 4). Second, to visualize the comprehensive contributions in a given voxel to both processes, we displayed Haufe *t*-values on the same brain slices using a two-dimensional colormap (Fig. 5A). The horizontal dimension represents *t*-values for pain increase vs. no change (green/red axis: deactivation/activation during pain increase), and vertical dimension reflects pain relief vs. no change (yellow/blue axis: deactivation/activation during relief). With this color coding, turquoise and orange map to regions with continuous increase-relief encoding (active in both but with opposite signs), whereas chartreuse (yellow-green) and purple denote regions with similar response in both conditions (having the same sign). Third, we extracted cluster-level Haufe *t*-values for both processes and visualized the condition-wise contributions on a scatter plot (Fig. 5B). Specifically, within PARES multivariate pattern (FDR *q* < .05), we extracted clusters (*K_E_* > 10 voxels). The dominant contribution of each cluster was determined by the signed maximal absolute Haufe *t*-values (peaks with extreme responses), yielding paired *t*-values that summarize cluster-level BOLD responses to pain increase and relief. These paired values were plotted on a scatter plot, where the *x*-axis represents *t*-values for pain increase vs. no change and the *y*-axis for relief vs. no change. The clusters were color coded using the same two-dimensional colormap described above. Together, these approaches provide a comprehensive picture of how the voxels/clusters in the PARES predictive pattern encode pain increase and relief.

In the Haufe-*t* maps, we observed a large cluster spanning the S1 and IPL with heterogeneous responses during pain increase and relief (Fig. 4C). To further disentangle the contributions within these two regions associated with distinct processes (i.e., nociception and modulation, respectively), each voxel in this cluster was assigned to the S1 or IPL by a forced choice using the probabilistic Juelich histological brain atlas (*17*) (resulting contours see Supplementary Fig. S5). Haufe-*t* values in the S1 and IPL were then visualized in the scatter plot (Fig. 5B).

### Support for applying the PARES in independent studies

In addition to making the entire analysis code for training and evaluating the signature available in the project repository (see Data and materials availability), we have implemented an easy-to-use Python routine for applying the PARES to new data, along with a step-by-step tutorial (https://git.uni-due.de/pnl-lab/pares-signature), to foster cumulative scientific efforts and translational applications.

## List of Supplementary Materials

Supplementary Methods Supplementary Results

Fig. S1. Univariate group-level brain activity in the cross-validated healthy controls (HC) sample.

Fig. S2. Behavioral ratings and predictive performance of the existing pain signatures in the external validated chronic back pain (CBP) sample.

Fig. S3. Multivariate predictive map of PARES.

Fig. S4. Wordcloud plots illustrating functional relevance of the Haufe-*t* maps.

Fig. S5. Haufe-weight maps in the S1 and IPL.

Fig. S6. FDR-significant regions in the PARES multivariate prediction.

Table S1. Metrics for testing the performance of multivariate predictive signatures.

Table S2. Clusters with positive predictive contributions surviving FDR correction (FDR *q* < .05; *K_E_* >= 5 voxels).

Table S3. Clusters with negative predictive contributions surviving FDR correction (FDR *q* < .05; *K_E_* >= 5 voxels).

References (*50*-*54*)

## Acknowledgments

We would like to thank Tor Wager for sharing the Neurologic Pain Signature (NPS), Giuseppe Gallitto for methodological support, and Evgenij Knorr for technical support. We also thank Folker Meyer and his colleagues at the Institute for Artificial Intelligence in Medicine (IKIM), Germany, for providing access to the high-performance computing cluster used for part of the analyses.

## Author contributions

Conceptualization: KS, KF, UB

Methodology: JL, BK, TS, UB

Investigation: JK, FS Supervision: TS, UB

Writing - original draft: JL, TS, KW, UB

Writing - review & editing: JL, TS, KW, UB, KS, KF, FS, LB, BK, JK

## Data and materials availability

The participant-level fMRI data used in this study will be available in the public repository upon acceptance of the manuscript.

The data and codes for reproducing the main analyses and figures, and a tutorial on applying PARES to new data are available at https://git.uni-due.de/pnl-lab/pares-signature.

