## supplements for "Beyond pain: a brain-based biomarker predicts individual pain relief"

[†] shared last authorship

### 1 Supplementary Methods

#### 1.1 Participants exclusion criterion

A total of 84 healthy participants (HC-sample) and 78 individuals with chronic back pain (CBP-sample) were recruited via advertisements. Exclusion criteria for both groups included age < 18 or > 75 years, severe neurological or psychiatric disorders, regular medication use (including hormonal contraception), MRI contraindications, pregnancy or breastfeeding, capsaicin allergy, dermatological abnormalities on the volar forearm, BMI < 18 or > 30, and left-handedness. For the healthy sample, a history of acute or chronic pain was an additional exclusion criterion. Patients were recruited through local advertisements and the Interdisciplinary Centre for Pain Medicine at University Hospital Essen. Eligibility was initially assessed by trained experimenters using a structured telephone screening and subsequently confirmed by pain-medicine physicians based on medical history and clinical examination. Patients were eligible if they had non-specific CBP, defined as remitting or persistent pain lasting >12 weeks in the absence of specific spinal pathology, nerve-root involvement, postsurgical or post-traumatic etiology, consistent with European CBP guidelines. Ongoing treatments had to remain stable for at least three weeks before the study. Patients receiving >100 mg morphine equivalents per day were excluded. Details of data exclusions are provided in *(14)*.

After applying these criteria and data-quality checks, n = 61 healthy participants (31 males, 30 females; mean age, 37.51 ± 13.28 years) and n = 58 individuals with CBP (25 males, 33 females; mean age, 39.00 ± 15.41 years) were included in the analyses, serving as the internal and external validation samples, respectively.

#### 1.2 Experimental procedure and experimental paradigm

The study was conducted on two experimental days. On day 1 (calibration day), participants underwent an individualized temperature calibration. Moderate continuous pain was elicited using a combined capsaicin and tonic heat pain model. Capsaicin cream (1%) was applied to a 3x3cm area on the forearm, enhancing thermal sensitivity and enabling low-level heat stimuli to maintain the heat pain levels with stable and lasting effects. After a 45min sensitization period during which participants completed self-report questionnaires, the cream was removed and thermal stimulation (Model ATS, Pathway System, Medoc, Israel) was applied to the capsaicin pre-treated area. Individual heat pain thresholds and temperature levels were then determined using a validated calibration procedure (for details see section 1.3).

On day 2 (experimental day), the capsaicin application and 45min sensitization were repeated, followed by reassessment of individual temperatures in the MRI scanner. Participants then completed the fMRI task, after which a high-resolution T1-weighted structural scan was acquired.

During the acquisition training of the conditioning paradigm, individually calibrated heat stimuli were paired with three visual geometric predictive cues for pain increase, relief and no-change conditions respectively. In this study, we focused on the behavioral pain intensity ratings and the brain activity during pain stimulations over 16 trials for each condition. Moderate heat pain was continuously applied to induce tonic pain baseline throughout the experimental phase. In the pain increase and relief trials, transient temperature increases and decreases were applied for 8s each, representing pain increase and relief respectively. A white fixation cross was displayed during the inter-trial-intervals for 6-11s. Following pain stimulations, pain intensity and (un)pleasantness ratings were collected on every fourth stimulus of the same type, yielding three ratings per condition and per question. Participants were given maximum 15s to rate each question, with a 0.5s interval in between. Paradigm details see *(14)*.

The software Presentation (www.neurobs.com) was used to present visual stimuli, trigger application of thermal stimuli and record behavioral data.

##### 1.3 Temperature calibration

To induce tonic heat pain while accounting for sensitization and habituation effects, each participant underwent a multi-step temperature calibration procedure on both experimental days. Pain intensity was rated using a 0–100 visual analog scale (VAS, 0 = "not painful at all", 100 = "unbearably painful"). This procedure identified three individualized temperature levels corresponding to intensely painful (VAS80), moderately painful (VAS40), and non-painful (VAS0) sensations, later used for pain increase, no change and pain relief conditions, respectively.

On day 1, calibration began with two staircase procedures starting at 28°C, increasing by 2°C (1°C per step when reaching 42°C, max 47°C) until participants reported ~VAS60. This temperature was then used in a following regression session, where ten temperature levels (−1.5°C to +3°C with a step of 0.5°C increment) were presented twice as an 8s heat stimulus before returning to baseline. After each heat stimulus, participants were asked to rate the pain intensity on the 0-100 VAS. Linear regression in R estimated temperatures corresponding to VAS80 and VAS60. The pain relief temperature level (VAS0) was defined as the temperature ~10°C below VAS60 (minimum 20°C) to elicit a relieving sensation from the tonic heat pain.

Because moderate tonic pain is particularly susceptible to habituation *(50)*, the temperature corresponding to VAS60 was chosen to later robustly induce a sustained pain intensity of VAS40 for the no-change condition. The subsequent training session started with the continuous application of the temperature corresponding to a pain intensity of VAS40, followed by three repetitions of temperature increases and decreases corresponding to the VAS80 andVAS0, presented in one of six counterbalanced orders. Participants rated pain intensity three times for each temperature change (increase, decrease) and five times for the no-change condition, to determine the robust temperature levels corresponding to VAS80, VAS0, and VAS40.

On day 2, the regression and training sessions were repeated inside the scanner to reassess the three temperature levels. The recalibrated temperatures were then used for the respective conditions during the main experiment.

##### 1.4 Time-course expression of PARES

To visualize how PARES captures pain dynamics during the time of pain stimulations, we extracted the PARES response from the participant-wise fMRI time courses during heat pain stimulations: pain increase, relief and no change. Specifically, for each participant, we regressed motion parameters from the preprocessed fMRI time courses per condition, analogous to a standard GLM nuisance-corrected analysis using TR-based beta-maps. Time-course expression of PARES were sampled from the stimulus onset through its following ~20s (9 TR). These event-aligned values were then aggregated across trials and participants to compute the mean response and 95% confidence intervals for each condition. The same procedure was applied in both samples.

##### 1.5 Examining correlations between PARES signature responses and head motion

To assess the potential influence of head motion on model predictions, we performed a quality control analysis using the first six components from anatomical component-based noise correction (aCompCor) *(44)* for each participant, derived by fMRIPrep. Both the mean and maximum absolute aCompCor values were computed across timepoints as motion metrics. Condition-wise PARES signature responses were calculated as the dot products between individual beta-maps and the mean of bootstrapped predictive weights. For each experimental condition, Pearson correlation was then calculated between the signature responses and both head motion metrics.

##### 1.6 Evaluating region-level feature importance in the PARES pattern

To complement the voxel-wise feature importance analysis reported in the main text, we conducted a region-level analysis of the PARES multivariate pattern. This step addresses a well-documented property of ridge regression models: because the L2 penalty distributes predictive weights across spatially extended, functionally homogeneous regions rather than concentrating them at individual voxels, contribution of a given voxel may be potentially underestimated *(49)*. Summarizing weights within anatomically defined regions therefore provides a more interpretable estimate of regional involvement in model prediction.

Following the same rationale and steps as in the voxel-level analysis, we summed bootstrapped voxel weights within predefined regions of interest (ROIs), using the MIST atlas (210 ROIs derived via a multi-level bootstrap analysis of stable clusters, BASC) *(51, 52)*, along with three additional brainstem subregions (periaqueductal gray, locus coeruleus and rostral ventromedial medulla) *(53, 54)*. This yielded region-level bootstrapped weights for 213 regions. Statistical inference followed the same framework as for voxel-level maps. Specifically, empirical one-tailed $p$-values were estimated for each region, and the tails of the bootstrap distributions were modelled using a GPD for regions passing a threshold of $p_{\text{one-tailed}} < .01$. Positive and negative weights were evaluated separately. Resulting $p$-values were corrected for multiple comparisons using false discovery rate (FDR $q < .05$), and significant regions indicated reliable positive or negative associations with changes in subjective ratings (Δratings).

#### 2 Supplementary results

##### 2.1 Univariate results

In the cross-validated HC-sample, we conducted univariate group-level analyses using one-sample *t*-tests for the comparisons: pain increase vs. no change, pain relief vs. no change and pain increase vs. pain relief. The results were consistent with our multivariate results and Haufe-weight maps (Fig. S1).

##### 2.2 NPS and SIIPS performance in the CBP-sample

The NPS performance in the CBP-sample closely replicated that observed in the HC-sample (Fig. S2B, Table 1). Specifically, the NPS showed significant within-participant condition-classification performance for pain increase ($t(57) = 8.528$, $p_{\text{one-tailed}} < .0001$), but not for pain relief ($t(57) = 3.264$, $p_{\text{one-tailed}} = .9989$). An exploratory two-tailed test for pain relief sensitivity revealed a “paradoxical” effect, pointing in the opposite direction than expected: the nociception-related NPS-score was higher during pain relief than no-change baseline ($p_{\text{two-tailed}} = .0024$). We found a similar pattern for the SIIPS (Fig. S2C, Table 1).

Together, these results reinforce the robustness of our observations in the HC-sample: the existing pain signatures reliably capture neural processes underlying pain increase but fail to characterize pain relief. Furthermore, it demonstrates that this dissociation can be replicated in a clinically relevant population.

##### 2.3 Correlations between PARES signature responses and head motion

Across all conditions and both motion metrics (mean and maximum aCompCor), no significant correlations were observed between signature responses and head motion (all $p$s > .05). This indicates that model predictions were not systematically associated with participant head movement, supporting that motion did not confound the PARES predictive weights.

##### 2.4 Region-level significant contributions to the PARES

At the region level, areas with significant positive contributions (FDR $q < .05$) were consistent with the voxel-level results, including the anterior insula, posterior insula, thalamus, cerebellum and visual area. Significant negative contributions were found in the orbitofrontal cortex, amygdala, intraparietal sulcus and inferior frontal gyrus (Fig. S6).

#### 3 Supplementary figures

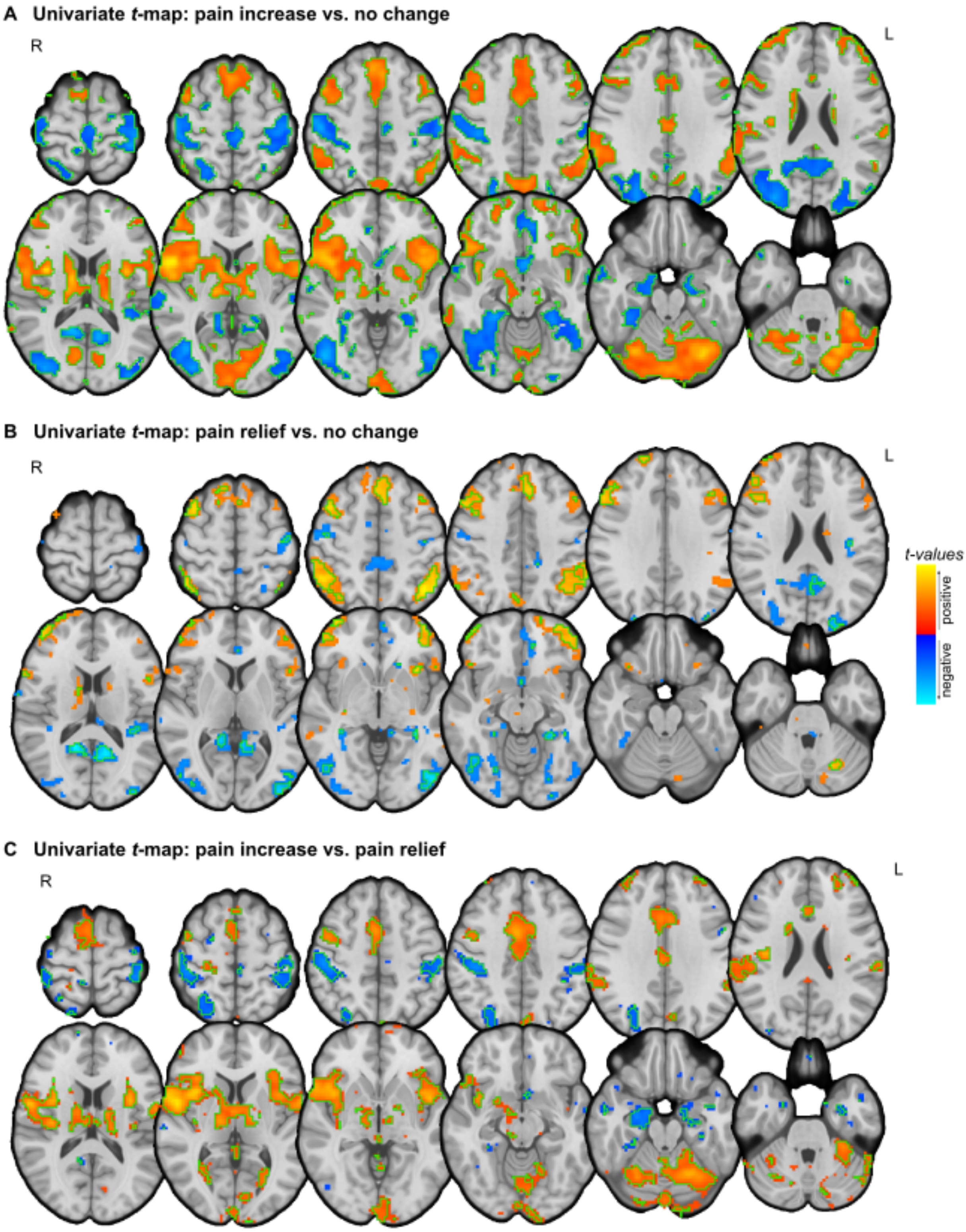


***Fig. S1. Univariate group-level brain activity in the cross-validated healthy controls (HC) sample.*** *Brain activations and deactivations for* ***(A)*** *pain increase vs. no change,* ***(B)*** *pain relief vs. no change and* ***(C)*** *pain increase vs. pain relief. Areas showing significant activity (one-sample t-tests,* $p_{uncorr} < .01$*) are depicted in warm colors (activation) and cool colors (deactivation) separately. Voxels in green contour are thresholded at FDR q < .05. L, left; R, right.*

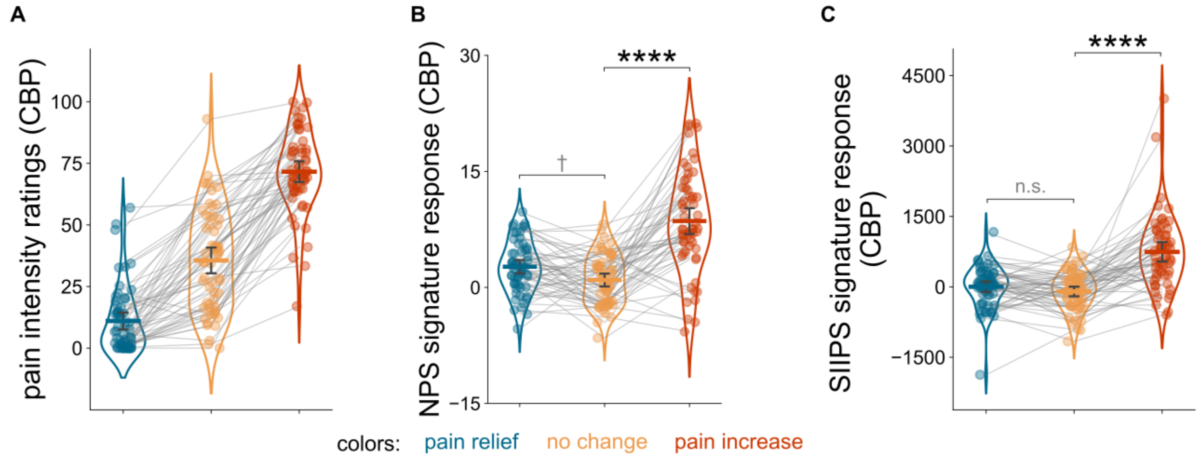


***Fig. S2. Behavioral ratings and predictive performance of the existing pain signatures in the external validated chronic back pain (CBP) sample.*** ***(A)*** *Behavioral pain intensity ratings during pain increase, pain relief and no change conditions in the CBP-sample. Predictive performance of the* ***(B)*** *Neurologic Pain Signature (NPS) and* ***(C)*** *Stimulus Intensity Independent Pain Signature (SIIPS). Both NPS and SIIPS significantly differentiated pain increase from no change but failed to distinguish pain relief from no change. Within-individual predictive performance was assessed using paired t-tests. Statistical significance was evaluated via permutation testing with 10,000 iterations. In all panels, violin plots show value distributions, with horizontal lines indicating means and error bars representing 95% bootstrap confidence intervals. Each dot represents one participant and gray lines connect within-participant conditions.* †*significant in the two-tailed tests but with an effect opposite to hypotheses;* ****$p < .0001$*; n.s., not significant.*

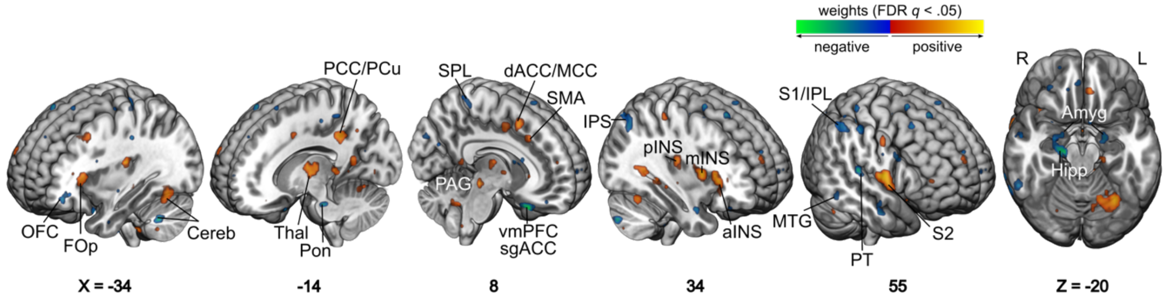


***Fig. S3. Multivariate predictive map of PARES.*** *Voxels with significant positive (warm colors) and negative (cool colors) predictive weights (bootstrap-derived FDR q < 0.05) indicate brain activity patterns that contributed to PARES model prediction. Positive weights were concentrated in pain-associated regions, including the insula, S2, thalamus, dACC/MCC, brainstem/PAG, cerebellum, and motor cortex, whereas negative weights involved regions linked to valuation, affect, and higher-order control, including the OFC, vmPFC, hippocampus, amygdala, IPS, SPL, and PCC/precuneus. FDR, false discovery rate; OFC, orbitofrontal cortex; FOp, frontal operculum; Cereb, cerebellum; Thal, thalamus; Pon, pons; PCC, posterior cingulate cortex; PCu, precuneus; PAG, periaqueductal gray; vmPFC, ventromedial prefrontal cortex; sgACC, subgenual anterior cingulate cortex; SPL, superior parietal lobule; dACC, dorsal anterior cingulate cortex; MCC, midcingulate cortex; SMA, supplementary motor area; IPS, intraparietal sulcus; pINS, posterior insula; mINS, mid-insula; aINS, anterior insula; MTG, middle temporal gyrus; PT, planum temporale; S2, secondary somatosensory cortex; S1, primary somatosensory cortex; IPL, inferior parietal lobule; Amyg, amygdala; Hipp, hippocampus; L, left; R, right.*

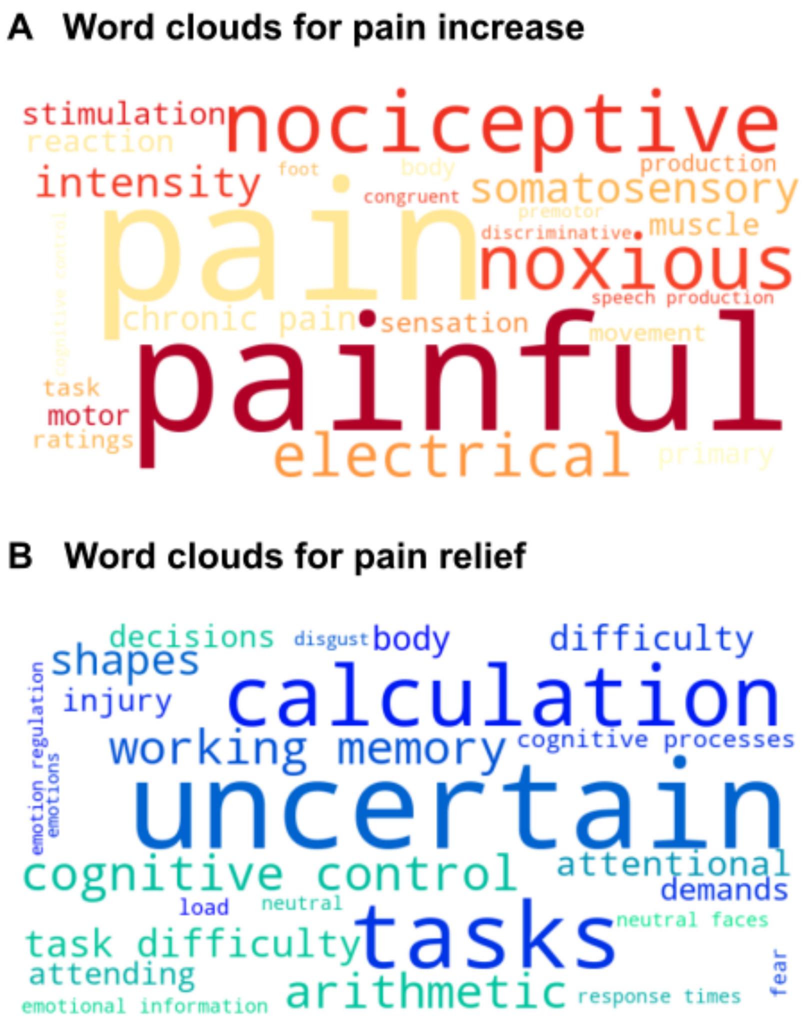


***Fig. S4. Wordcloud plots illustrating functional relevance of the Haufe-t maps.*** *Functional items associated with the Haufe-t maps of pain increase and pain relief were derived from the study pool of Neurosynth. Items and their correlation coefficients were extracted and visualized as the word clouds for* ***(A)*** *pain increase and* ***(B)*** *pain relief, with font size representing the strength of associations in Neurosynth.*

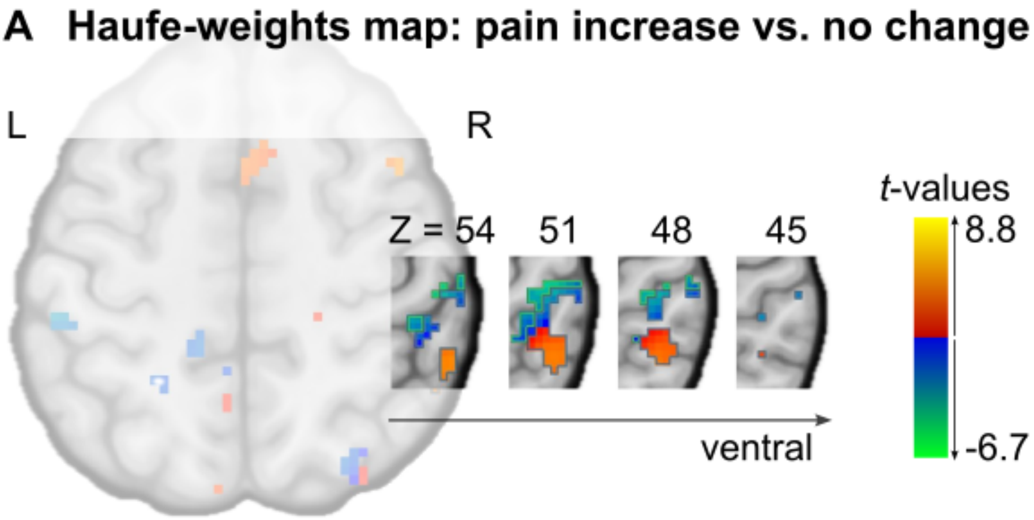


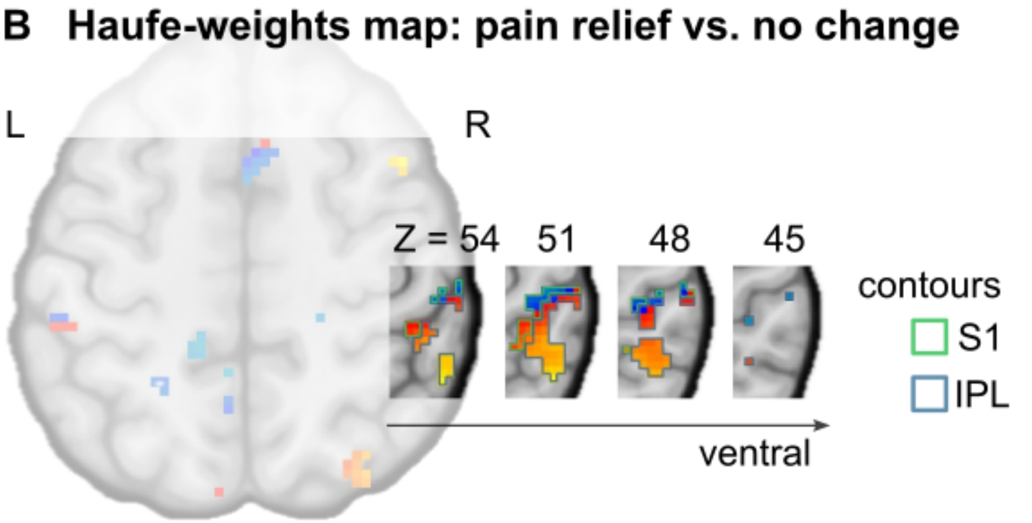


***Fig. S5. Haufe-weight maps in the S1 and IPL.*** *Contours display the primary somatosensory cortex (S1) and inferior parietal lobule (IPL) regions based on the probabilistic Juelich histological brain atlas (17), overlaid on the Haufe-t maps for* ***(A)*** *pain increase and* ***(B)*** *pain relief. The same t-value range is used for the pain-increase and pain-relief maps to facilitate direct visual comparison. L, left; R, right.*

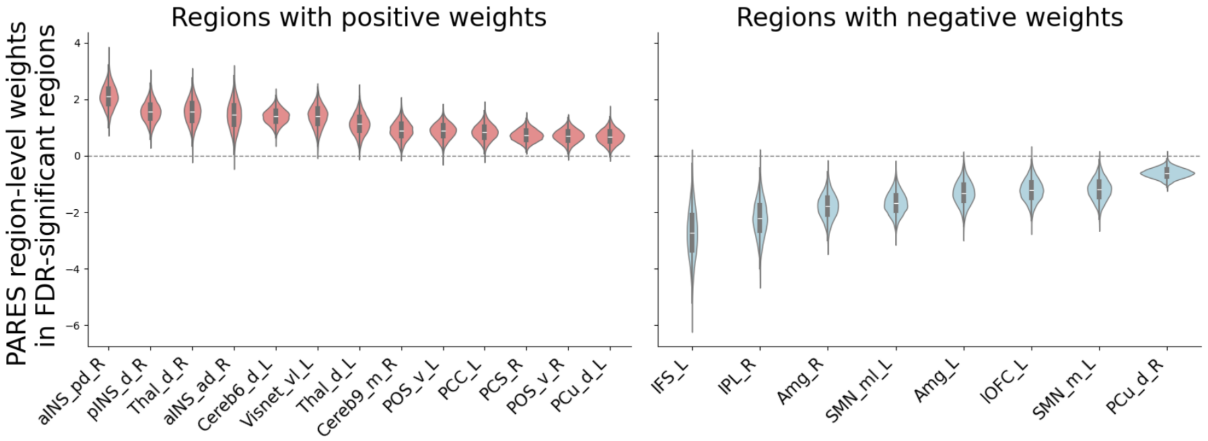


***Fig. S6. FDR-significant regions in the PARES multivariate prediction.*** *Violin plots show bootstrapped predictive weights for the significant regions (FDR q < .05) in the region-level feature importance analysis (bootstrapping test with 10,000 iterations). As described in the Supplementary Methods 2.4, the voxel-level weights in the PARES predictive pattern were summed within 213 anatomical regions (210 MIST ROIs and three brainstem subregions from (53, 54). Regions with positive and negative significant weights are displayed in the left and right panels, respectively. aINS_pd_R, right anterior insula (posterior dorsal); pINS_d_R, right dorsal posterior insula; Thal_d_R, right dorsal thalamus; aINS_ad_R, right anterior insula (anterior dorsal); Cereb6_d_L, left cerebellum VI (dorsal); Visnet_vl_L, left visual network (ventrolateral); Thal_d_L, left dorsal thalamus; Cereb9_m_R, right cerebellum IX (medial); POS_v_L, left parietal occipital sulcus (ventral); PCC_L, left posterior cingulate cortex; PCS_R, right paracentral sulcus; POS_v_R, right parietal occipital sulcus (ventral); PCu_d_L, left dorsal precuneus; IFS_L, left inferior frontal sulcus; IPL_R, right inferior parietal lobule; Amyg_R, right amygdala; SMN_ml_L, left sensorimotor network (medial); Amyg_L, left amygdala; lOFC_L, left lateral orbitofrontal cortex; SMN_m_R, right sensorimotor network (medial); PCu_d_R, right dorsal precuneus.*

#### 4 Supplementary tables

Table S1. Metrics for testing the performance of multivariate predictive signatures.

Table S2. Clusters with positive predictive contributions surviving FDR correction (FDR $q < .05$; $K_E >= 5$ voxels).

Table S3. Clusters with negative predictive contributions surviving FDR correction (FDR $q < .05$; $K_E >= 5$ voxels).

***Table S1. Metrics for testing the performance of multivariate predictive signatures.***

| Performance metric | Test statistic | Signatures tested |
|---|---|---|
| P1. Pooled inter-individual performance | Pearson correlation between observed and predicted behavior, across conditions and participants | NPS, SIIPS, PARES |
| P2. Pooled intra-individual performance | Mean of the within-participant Pearson correlation between observed and predicted behavior, across conditions | NPS, SIIPS, PARES |
| P3. Sensitivity to pain increase | Paired *t*-test between the "pain increase" and "no change" conditions | NPS, SIIPS, PARES |
| P4. Sensitivity to pain relief | Paired *t*-test between the "pain relief" and "no change" conditions | NPS, SIIPS, PARES |
| P5. Sign-based classification accuracy | Proportion of correct predictions (accuracy), when using the signs of the predictions to distinguish the pain increase and relief conditions | PARES |

***Table S2. Clusters with positive predictive contributions surviving FDR correction (FDR q < .05; $K_E$ >= 5 voxels).*** *abs., absolute; PAG, periaqueductal gray; Thal_p, posterior thalamus; Hipp, hippocampus; S2, secondary somatosensory cortex; pINS, posterior insula; PreCG, precentral gyrus; Cereb, cerebellum; mINS, mid-insula; aINS, anterior insula; Thal_a, anterior thalamus; SMA, supplementary motor area; dACC, dorsal anterior cingulate cortex; PCC, posterior cingulate cortex; PCu, precuneus; PT, planum temporale; FOp, frontal operculum; M1, primary motor cortex; MTG, middle temporal gyrus; vlPFC, ventrolateral prefrontal cortex; MFG, middle frontal gyrus; IPL, inferior parietal lobule; lOC, lateral occipital complex; V3, third visual cortex.*

| Cluster size ($mm^3$) | X | Y | Z | Region | Multivariate weight (peak) | FDR *q* (peak) | increase | relief |
|---|---|---|---|---|---|---|---|---|
| | | | | | | | Signed max abs. Haufe-*t* | Signed max abs. Haufe-*t* |
| 86 | 6 | -28 | -11 | PAG | 0.0179 | < .0001 | 2.528 | -1.516 |
| 954 | 18 | -23 | 8 | Thal_p | 0.0176 | < .0001 | 6.862 | -1.424 |
| 138 | 35 | -38 | -3 | Hipp | 0.0175 | < .0001 | -3.372 | -4.435 |
| 2931 | 55 | 2 | 11 | S2 | 0.0171 | 0.0001 | 8.679 | -2.679 |
| 659 | 35 | -16 | 17 | pINS | 0.0171 | < .0001 | 6.620 | -2.310 |
| 676 | -37 | -18 | 17 | pINS | 0.0165 | < .0001 | 3.693 | -3.836 |
| 1960 | -59 | -3 | 14 | PreCG | 0.0162 | < .0001 | 6.213 | -2.640 |
| 1890 | -29 | -63 | -17 | Cereb | 0.0152 | < .0001 | 6.460 | -3.167 |
| 659 | -14 | -8 | 11 | Thal | 0.0147 | 0.0004 | 6.295 | -1.173 |
| 971 | 33 | 9 | 8 | mINS | 0.0147 | < .0001 | 8.810 | -1.947 |
| 1248 | 30 | 24 | 8 | aINS | 0.0145 | 0.0073 | 6.027 | 2.807 |
| 104 | 11 | -3 | 5 | Thal_a | 0.0144 | 0.0014 | 5.618 | 1.353 |
| 225 | 13 | -43 | -51 | brainstem | 0.0131 | < .0001 | 2.320 | -3.220 |
| 1214 | 8 | 14 | 50 | SMA | 0.0125 | 0.0044 | 5.281 | -2.643 |
| 190 | 11 | 24 | 39 | dACC | 0.0121 | 0.0188 | 4.349 | -1.015 |
| 294 | -4 | -48 | 3 | PCC | 0.0118 | 0.0004 | 2.875 | -4.582 |
| 104 | 18 | -63 | 36 | PCu | 0.0117 | 0.0044 | 3.190 | 1.112 |
| 104 | -47 | 19 | -23 | PT | 0.0117 | 0.0064 | 2.152 | -0.373 |
| 728 | -29 | 24 | 11 | FOp | 0.0116 | 0.0011 | 6.166 | 1.179 |
| 329 | 30 | -58 | -23 | Cereb | 0.0114 | < .0001 | 5.473 | -2.252 |
| 572 | 48 | -3 | 42 | PreCG | 0.0114 | < .0001 | 4.574 | -2.205 |
| 208 | 35 | -30 | 56 | M1 | 0.0112 | < .0001 | -2.015 | -2.660 |
| 86 | -64 | -40 | -14 | MTG | 0.0105 | 0.0222 | 2.375 | 0.753 |
| 86 | 38 | 39 | 5 | vlPFC | 0.0104 | 0.0448 | 5.133 | 2.699 |
| 797 | -27 | 24 | 39 | MFG | 0.0099 | 0.0241 | 2.543 | 1.998 |
| 86 | -57 | -48 | 31 | IPL | 0.0096 | 0.0079 | 3.836 | 1.829 |
| 173 | 25 | -88 | 5 | lOC | 0.0095 | 0.0018 | -2.811 | -3.581 |
| 225 | -4 | -73 | 39 | PCu | 0.0088 | 0.0152 | 5.401 | 2.159 |
| 104 | 23 | -73 | -14 | V3 | 0.0077 | 0.0029 | 1.487 | -2.922 |

***Table S3. Clusters with negative predictive contributions surviving FDR correction (FDR q < .05; $K_E$ >= 5 voxels).*** *abs., absolute; vmPFC, ventromedial prefrontal cortex; Thal, thalamus; Hipp, hippocampus; LG, lingual gyrus; OFC, orbitofrontal cortex; PT, planum temporale; SMG, supramarginal gyrus; Amyg, amygdala; SMG, supramarginal gyrus; sgACC, subgenual anterior cingulate cortex; S1, primary somatosensory cortex; M1, primary motor cortex; IPS, intraparietal sulcus; SFG, superior frontal gyrus; IPL, inferior parietal lobule; MTG, middle temporal gyrus; V2, secondary visual cortex; IFG, inferior frontal gyrus.*

| **Cluster size (mm³)** | **X** | **Y** | **Z** | **Region** | **Multivariate weight (peak)** | **FDR *q* (peak)** | **increase** | **relief** |
|---|---|---|---|---|---|---|---|---|
| | | | | | | | **Signed max abs. Haufe-*t*** | **Signed max abs. Haufe-*t*** |
| 485 | 8 | 24 | -28 | vmPFC | -0.0176 | < .0001 | -2.050 | 1.393 |
| 104 | 1 | -25 | 0 | Thal | -0.0162 | 0.0200 | -0.908 | 1.847 |
| 1127 | 18 | -20 | -20 | Hipp | -0.0155 | 0.0002 | -5.433 | -1.438 |
| 86 | -12 | -25 | -28 | Pons | -0.0138 | 0.0087 | -0.799 | 1.660 |
| 86 | -37 | -1 | -34 | fusiform | -0.0133 | 0.0327 | -2.436 | 1.089 |
| 208 | 16 | -75 | 0 | LG | -0.0131 | 0.0003 | -1.966 | -1.359 |
| 242 | -34 | 44 | -6 | OFC | -0.0125 | 0.0214 | 2.209 | 4.212 |
| 277 | 53 | -28 | 11 | PT | -0.0122 | 0.0140 | -2.496 | -1.043 |
| 104 | 28 | 2 | -25 | Amyg | -0.0121 | 0.0429 | -3.050 | -0.660 |
| 173 | 38 | -25 | 39 | SMG | -0.0120 | 0.0004 | -5.506 | -0.824 |
| 988 | -44 | -33 | 39 | SMG | -0.0119 | 0.0112 | -3.947 | 3.350 |
| 294 | -19 | -6 | -23 | Amyg | -0.0118 | < .0001 | -4.637 | -1.663 |
| 208 | 13 | 32 | -9 | sgACC | -0.0117 | 0.0060 | -2.418 | 1.690 |
| 646 | 60 | -23 | 50 | S1 | -0.0114 | 0.0004 | -6.752 | 2.599 |
| 416 | -29 | -25 | 67 | M1 | -0.0111 | 0.0054 | -4.254 | -2.428 |
| 208 | -39 | -48 | 33 | IPS | -0.0107 | 0.0001 | 1.009 | 3.098 |
| 346 | 23 | 37 | 28 | SFG | -0.0106 | 0.0004 | 1.318 | 2.419 |
| 1343 | 60 | -25 | 50 | IPL | -0.0105 | 0.0004 | -5.671 | 5.256 |
| 208 | -52 | -23 | 33 | S1 | -0.0105 | 0.0082 | -1.806 | 1.645 |
| 693 | 65 | -50 | -9 | MTG | -0.0104 | 0.0456 | 2.180 | 2.503 |
| 138 | -49 | 34 | 17 | IFG | -0.0102 | 0.0248 | 1.960 | 2.680 |
| 294 | -54 | -28 | 8 | PT | -0.0102 | 0.0161 | -3.483 | -2.541 |
| 468 | 13 | -83 | 19 | V2 | -0.0100 | 0.0175 | -1.712 | 1.183 |
| 641 | 33 | -68 | 42 | IPS | -0.0093 | 0.0083 | -2.422 | 3.657 |
| 104 | 55 | -6 | -20 | MTG | -0.0092 | 0.0302 | -4.734 | -1.710 |
